# Strategies for increasing the depth and throughput of protein analysis by plexDIA

**DOI:** 10.1101/2022.11.05.515287

**Authors:** Jason Derks, Nikolai Slavov

## Abstract

Accurate protein quantification is key to identifying protein markers, regulatory relationships between proteins, and pathophysiological mechanisms. Realizing this potential requires sensitive and deep protein analysis of a large number of samples. Toward this goal, proteomics throughput can be increased by parallelizing the analysis of both precursors and samples using multiplexed data independent acquisition (DIA) implemented by the plexDIA framework: https://plexDIA.slavovlab.net. Here we demonstrate the improved precisions of RT estimates within plexDIA and how this enables more accurate protein quantification. plexDIA has demonstrated multiplicative gains in throughput, and these gains may be substantially amplified by improving the multiplexing reagents, data acquisition and interpretation. We discuss future directions for advancing plexDIA, which include engineering optimized mass-tags for high-plexDIA, introducing isotopologous carriers, and developing algorithms that utilize the regular structures of plexDIA data to improve sensitivity, proteome coverage and quantitative accuracy. These advances in plexDIA will increase the throughput of functional proteomic assays, including quantifying protein conformations, turnover dynamics, modifications states and activities. The sensitivity of these assays will extend to single-cell analysis, thus enabling functional single-cell protein analysis.

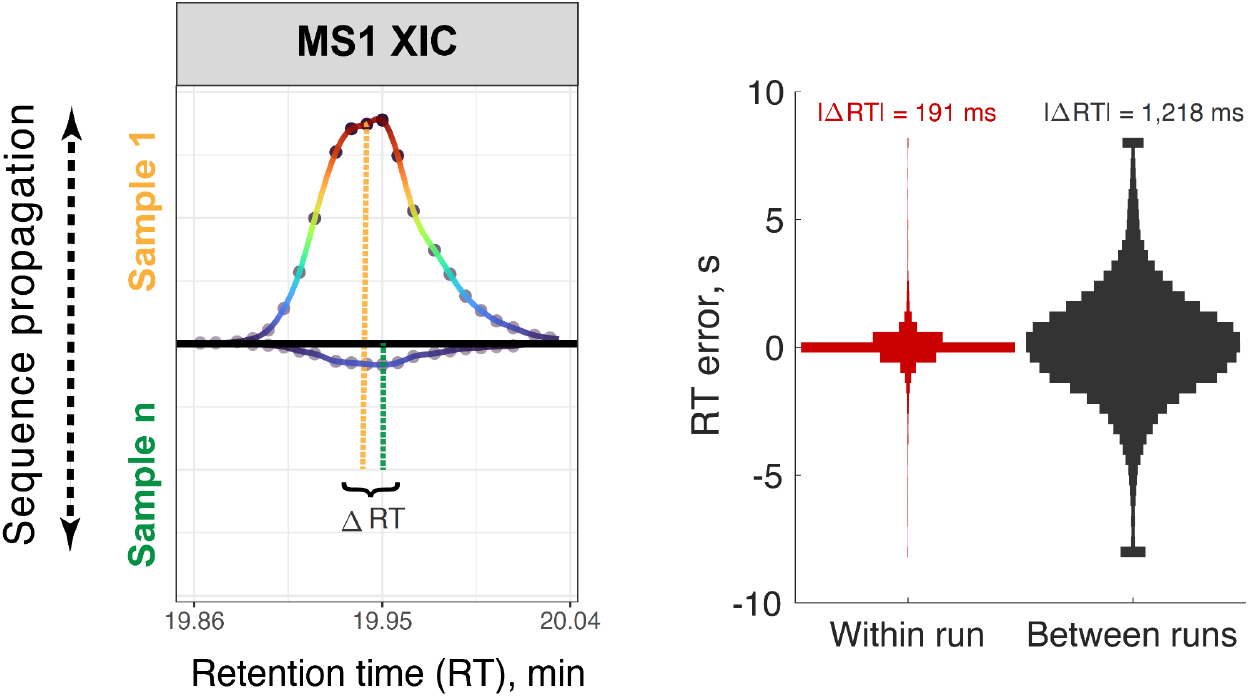

## Introduction

Tandem mass-spectrometry (MS) has long been established as the most specific, comprehensive, and versatile method for protein analysis^1–3^. However, the sensitivity and throughput of MS have traditionally limited the biomedical applications of MS proteomics. These limitations are increasingly mitigated by new approaches that increase the sensitivity^4–6^ and throughput^7,8^ of MS-based proteomics. Many of these advances take advantage of data independent acquisition (DIA), which was introduced decades ago^9^ and has developed into powerful methodologies^10–15^.

In this *Perspective*, we focus on one approach for increasing both the sensitivity and throughput of MS-based protein analysis: plexDIA^16^. The wide isolation windows used by DIA allow for parallel accumulation of ions for fragmentations and MS2 analysis, which may enable analyzing many peptides using the long ion accumulation times required for single-cell proteomics^17^. Indeed, DIA allows obtaining MS2 fragmentation spectra from all detectable peptide features even when using long ion accumulation times, as shown in Fig. 1a. This makes it attractive for analyzing small samples, such as single cells^18^. However, long accumulation times reduce the number of times elution peaks are sampled (Fig. 1b). Furthermore, wide isolation windows increase ion interference. These factors raise challenges both for sequence identification and protein quantification. The challenges may be partially mitigated, e.g., by introducing multiple MS1 survey scans for increasing the points per peak (Fig. 1b)^16,19^, and by other approaches outlined in this *Perspective*.

**Figure 1:**
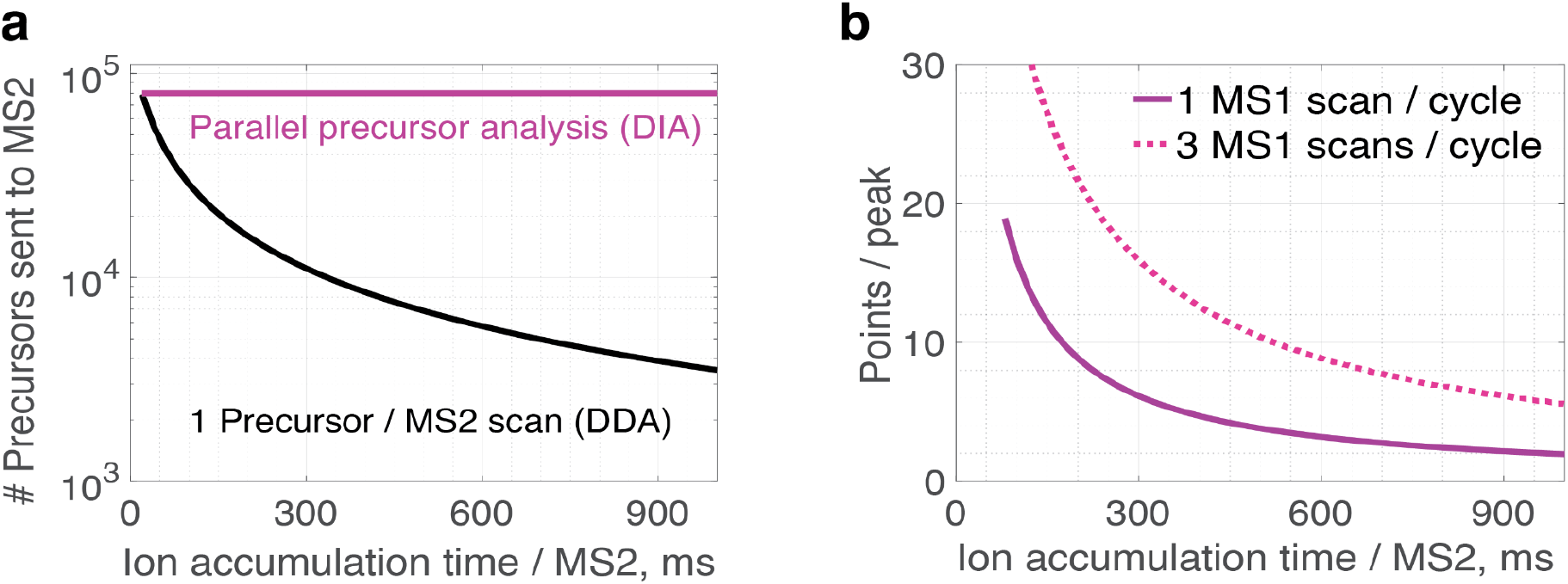
Parallel precursor isolation and fragmentation enable analyzing all detectable precursors even when using long ion accumulation times for MS2 scans. **a**, As ion accumulation times increase, the number of precursors that can be fragmented and analyzed at MS2 level decreases for data dependent acquisition (DDA) analysis. The DDA graphs show a theoretical estimate for the maximum number of precursors that can be analyzed as a function of ion accumulation times for MS2 scans while using a 60 min active gradient and assuming full duty cycles^18^. In contrast, parallel isolation and fragmentation of precursors by DIA allows for analyzing all detectable precursors even when using long ion accumulation times for MS2 scans. **b**, As accumulation times increase, the points per elution peak decrease as illustrated for duty cycles having 10 MS2 scans per cycle and either 1 or 3 MS2 scans per duty cycle. Elution peaks were modeled as 20 seconds at base for 60 min active gradient, and narrower peaks will have fewer sampling points.

MS analysis of a single human cell detects over 60,000 peptide-like precursors^20^ Parallel isolation and fragmentation of the precursors may allow analyzing all of them at the MS2 level(Fig. 1), but it does not guarantee identification, accurate quantification and high throughput. Achieving these goals can benefit from sample multiplexing and algorithms for enhanced sequence identification and quantification, thus creating exciting technological and methodological opportunities^18^. We discuss such outstanding opportunities for major gains that may be enabled by optimized mass tags and algorithms for peptide sequence identification and quantification.

### Development of multiplexed data independent acquisition

Sample multiplexing by data independent acquisition (DIA) was demonstrated by Minogue *et al.^21^* using metabolic labeling by Stable Isotope Labeling with Amino acids in Cell culture (SILAC). Subsequently, it was extended to pulsed SILAC^22,23^, which allowed measuring protein turnover rates. These studies convincingly demonstrated that the complex spectra of multiplexed DIA can be interpreted and used to quantify proteins. Yet, this came at a price: The number of proteins quantified by label-free DIA (LF-DIA) was about 2-fold larger than the number of proteins quantified by pulsed SILAC DIA^23^. Furthermore, the effect of metabolic multiplexing on quantitative accuracy was not directly benchmarked, and subsequent studies highlighted the challenges for quantification^24,25^.

More recently, Pino *et al.^26^* rigorously benchmarked quantitative accuracy by SILAC-DIA and found that it exceeds the accuracy achieved by SILAC-DDA. Yet, SILAC-DIA quantified about 2-fold fewer peptide sequences in the mixed heavy and light samples compared to only heavy or only light samples. This result is similar to the 2-fold reduced proteome coverage reported by Liu *et al.^23^* and reinforced the notion that multiplexing with DIA may not increase throughput (defined as the number of quantitative data points per unit time) over label-free DIA. Furthermore, the throughput of SILAC-DIA did not exceed the throughput of SILAC-DDA. Indeed, the numbers of precursors quantified in the mixed heavy and light samples were very similar for SILAC-DDA and SILAC-DIA, suggesting that SILAC-DIA did not increase the throughput and proteome coverage compared to SILAC-DDA^26^. These results highlight both the potential and the challenges of multiplexed DIA. Indeed, multiple implementations of multiplexed DIA have reported proteome coverage of about 2,000 proteins or fewer, significantly below the proteome coverage that may be achieved by the corresponding LF-DIA analysis^21,27–29^. This reduction in proteome coverage by multiplexed DIA reduced its appeal despite its demonstrated ability to multiplex samples.

#### Mass tags for multiplexed DIA

Multiplexed DIA can be implemented with different types of mass tags, each type having their own distinct characteristics. One important property that distinguishes mass tags is whether labeled peptides produce sample-specific precursors and fragments, as listed in Table 1 along with representative examples of tags. This property determines the ability to support quantification and sequence identification using MS1 and/or MS2 level measurements as explained below.

**Table 1.** Types of amine-reactive mass tags that can multiplex samples for DIA analysis. Mass tags are classified based on their ability to generate sample-specific precursors and fragments. ^a^For peptides cleaved after lysine (e.g., when proteins are digested with lys-C), both b and y ions are sample-specific. For other peptides, only b ions are sample-specific while y ions may not be sample-specific. ^b^Upon fragmentation, peptides labeled with isobaric mass tags produce reporter ions (RI) that are not peptide and sample specific and cannot support peptide quantification^35^. They also produce complement RI (sample specific tag fragments attached to peptide fragments) that may be peptide and sample specific^34^. A subset of these complement RIs with non-overlapping isotopic envelopes may support peptide and sample specific quantification at the MS2 level^34,35^.

| Mass tag type | Sample-specific precursors | Sample-specific fragments | Examples |
| --- | --- | --- | --- |
| Type I | Yes | Yes<sup>a</sup> | mTRAQ, Dimethyl<sup>32</sup>, TMT0/TMT/TMTsh, mdDiLeu<sup>33</sup> |
| Type II | Yes | No | Ac-IP<sup>29</sup> |
| Type III | No | Yes<sup>b</sup> | TMTc<sup>34</sup> |

Type I mass tags result in sample-specific precursors and fragments and thus enable quantitation and sequence identification at both MS1 and MS2-level, Table 1. This specificity maximizes the confidence of identifying the composition of each sample^16^ and provides a reliability estimate based on the consistency of MS1 and MS2 level quantification^4^. These benefits come at the expense of more complex MS2 spectra. These mass tags can include neutron-encoded (NeuCode) chemical labels that introduce small mass-offsets due to the mass defect of neutron binding energy^21,27,30,31^. Depending on the resolution of MS analysis, such NeuCode labels may appear as isobaric (at low resolution) or as non-isobaric (at high resolution); thus, if fragments can be analyzed with low resolving power as isobaric and precursors analyzed with high resolving power as non-isobaric, these tags will function as Type II mass tags (Table 1). Using the mass defect to introduce sub-Dalton mass offsets offers the possibility of achieving high-plexDIA. However, so far methods implementing such mass tags have required long orbitrap scan times, which slows the duty cycles and reduces proteome coverage.

Type II mass tags result in sample-specific precursors and fragments that are shared across samples. Thus, these tags have less complex MS2 spectra. However, the absence of sample-specific fragments sacrifices MS2 evidence for the peptides present in each sample, and thus limits the specificity of sequence identification. Furthermore, these tags do not support MS2-level quantification needed for the quantification consistency estimates possible with Type II tags^4,16^.

Type III mass tags are isobaric and result in precursors that are shared across samples and some fragments that may be sample and peptide specific. Only complement reporter ions attached to peptide specific fragments may be sample and peptide specific^34^. Thus, only a subset of the fragments may be used for sample-specific peptide identification and quantification^35^. When using wide isolation windows (as commonly done with DIA), avoiding overlap between the isotopic envelopes of complement reporter ions requires using only tags having reporter ions that are separated by at least 4Da or by detectable mass defect. This requirement means that only a small subset of the TMT tags can be used together for multiplexing. These limitations of Type III mass tags informed our choice to use non-isobaric mass tags for plexDIA^16,36^.

##### Increasing proteome coverage and accuracy with plexDIA

As discussed above, the complex spectra of multiplexed DIA have posed formidable challenges, especially to matching the proteome coverage of LF-DIA^21,23–29^. Towards overcoming these challenges and increasing the proteome coverage, we introduced plexDIA^16,37^. It uses Type I mass tags and a computational framework that allows increased throughput without sacrificing proteome coverage. Furthermore, plexDIA increased data completeness and the accuracy of relative protein quantification as benchmarked by mixed species proteomes.

plexDIA improved data-completeness by allowing consistent quantification of more proteins across diverse samples than what can be achieved with LF-DIA using 3-times less instrument time^16^. These gains stem from computational approaches leveraging the fact that isotopically labeled samples co-elute. Thus, peptide sequences which are confidently identified in one isotopic channel may be confidently propagated to other co-eluting channels because of the ability to accurately and precisely predict the m/z and retention time of each precursor and its corresponding fragments.

Achieving high quantitative accuracy when using Type I mass tags may be challenging since they increase the spectral complexity linearly with plex; therefore, multiplexed DIA has the potential to be affected by increased interference, which may result in reduced quantitative accuracy. Despite this potential, the accuracy of 3-plexDIA was made comparable to LF-DIA by limiting the impact of interferences through 1) quantifying peptides based on MS scans nearer the elution peak apex, and 2) developing an algorithm that quantifies precursors within a set relative to the most confidently assigned channel, as illustrated in Fig 2a. Both approaches are motivated by the principle that quantitation derived from the elution peak apex provides the strongest signal relative to interferences. Results from the bulk mixed-species plexDIA dataset are shown in Fig. 2b to assess the improvement of accuracy by the ‘translated quantification’ algorithm. While the algorithm does not improve MS1-level quantitation, translated MS2 quantities are more accurate than non-translated quantities as shown by smaller ratio errors.

**Figure 2:**
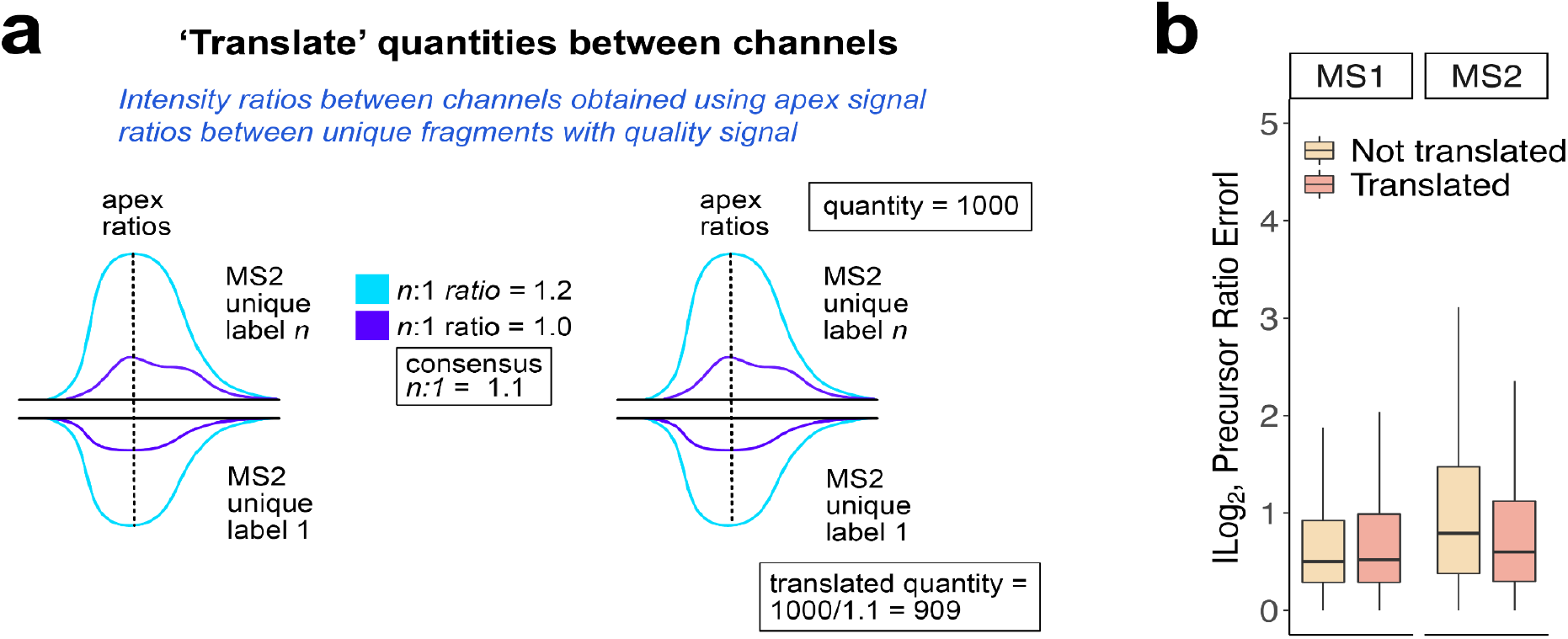
The accuracy of protein quantification at MS2-level increases with the translation algorithm. **a**, plexDIA uses a translation algorithm to reduce the impact of interferences by scaling the apexes of fragments from propagated sequences to the most confident sequence^16^. The algorithm uses the average fragment ratio to scale the quantity of the best quantified precursors to other precursors with the same sequence. This panel was adapted from Derks *et al.^16^*. **b**, Mixed species samples used to benchmark plexDIA performance^16^ were used to assess quantitative accuracy with and without translation. Boxplot distributions of MS1 and MS2-level deviations from the expected ratios are plotted for the precursor ratios (*n*=37,907) that were quantified in common across all samples. MS2-level quantitative accuracy improves in the “MS2 translated” condition (orange) relative to non-translated quantities (yellow).

MS2-level translation likely benefits from averaging ratios across many fragments as opposed to MS1-level which produces just a single apex ratio, which may explain the discrepancy.

## Opportunities for advancing plexDIA

While the demonstrated performance of plexDIA^16^ already provides substantial advantages for practical applications, it has much potential for further improvement^36^. plexDIA opens new avenues for methodological advances, both for developing new mass-tags for multiplexing and for advancing the computational frameworks for data interpretation. These developments are discussed in the subsections below, along with their interdependence and requirements towards MS instrumentation.

### Developing mass-tags optimized for plexDIA

Mass tags optimized for plexDIA can increase both the proteome coverage (by enhancing amino acid sequence identification) and the number of samples analyzed simultaneously (by increasing the plex), as discussed below.

### Optimizing mass tags for sequence identification

The fragmentation properties of mass tags are crucial for their utility for plexDIA. The desired fragmentation properties for optimal performance differ depending on the type of mass tags listed in Table 1: While Type I tags should be engineered to minimize fragmentation, Type II and III tags should be engineered to maximize fragmentation. For all types of tags, reducing spectral complexity and increasing sensitivity benefits from mass tags that do not generate fragments that are neither peptide specific nor sample specific. The mass tags used to benchmark plexDIA, mTRAQ, have the chemical structure of the isobaric iTRAQ and produce reporter ions as well. These reporter ions can be deleterious to peptide identification and quantification at MS2-level as they reduce a pool of fragments lacking peptide specificity. These non-specific fragments use up the limited capacity of ion traps and detectors without contributing to sample and peptide specific analysis. Thus, developing optimized mass-tags for plexDIA should limit undesirable fragmentation. Such tags should improve peptide identification and quantification by plexDIA.

Mass tags may be engineered to contribute additional benefits to sequence identifications and sensitive quantification. For example, mass tags may stabilize charge on fragment ions and thus increase the detectable fragments. plexDIA already benefited from the propensity of mTRAQ to stabilize b-ions, but this propensity can be further enhanced in the next generations of mass tags. As another benefit, mass tags can be engineered to contribute additional charge (such as by adding amine groups), which will increase the sensitivity of detection by MS. Such high-charge designs will be particularly useful for single-cell proteomics^35^. Another potential benefit for sensitive proteomics could be the increased signal from pooling peptide fragments originating from different samples. Such pooling happens when the same peptide sequence labeled with different mass tags from Type II generates the same fragments. This can also happen with the y-ions of Type I mass tags. Such pooling can enhance peptide sequence identification analogously to the pooling that happens with isobaric carriers^20,38^, but it also may limit the specificity of sample-specific sequence identification. Thus, rigorous models of amino acid sequence identification should ensure robust FDR estimations and benchmark them with mixed species experiments as described in the community white paper on single-cell proteomics^4^.

### Increasing the number of multiplexed samples

The multiplicative scaling of throughput by plexDIA has been demonstrated with a 3-plexDIA^16^, and we expect this framework to extend to suitably engineered higher plex mass tags^36,39^. The mass tags may be designed with both large (4Da or more) and small (mass defect sub-Dalton) mass shifts, similar to the design by the Coon laboratory^30^. The sub-Dalton differences should be large enough to be resolved without requiring MS scan times that would extend the time of duty cycles beyond the times that are optimal for maximizing sequence coverage. Designing such tags within the constraints required for optimal performance of plexDIA is challenging, but it has the potential to extend the multiplicative scaling of throughput high-plexDIA.

Realizing this potential also requires MS instrumentation and experimental designs that can keep up with the increasing complexity of MS spectra. The proof of principle plexDIA demonstration used Type I mass tags with 4Da mass-shifts, which increase the ion complexity at both MS1 and MS2 levels. Despite the added complexity, the 3-plexDIA quantified over 200,000 precursors (~8,000 protein groups per sample) while using 1-hour active gradient and a first-generation Q-Exactive. This analysis resulted in comparable quantitative accuracy to matched LF-DIA^16^, suggesting that the capacity of ion traps and detectors was not saturated by 200,000 precursors. Thus, we expect the possibility to further increase the number of accurately quantified precursors, especially when using optimized experimental designs with smaller isolation windows and newer instruments, such as high-field orbitraps or fast TOFs combined with narrow MS2 isolation windows. This potential for scaling is particularly great for single-cell proteomics because current methods quantify fewer precursors per single-cell sample. The current coverage of our 10,000 precursors quantified per single cell and 200,000 precursors quantified by Derks *et al.^16^* in a single run correspond to a 20-plex single-cell set. The analysis of such a set will be further facilitated by its smaller dynamic range, which reduces the potential for interference. Using ion mobility and/or higher resolution MS detectors can further increase the number of samples that can be multiplexed without undermining quantification accuracy. Therefore, we expect high-plexDIA to be particularly powerful in scaling up single-cell proteomics^39^.

### plexDIA algorithms for enhanced sequence identification

As discussed above, the plexDIA framework can allow propagating amino acid sequences within a run with much higher sensitivity and rigor than propagating sequences between runs. These capabilities stem from the co-elution of peptides labeled with non-isobaric isotopologous mass tags. Algorithms that effectively leverage the time information inherent in coeluting peptides can further amplify the power of sensitive and rigorous sequence propagation within plexDIA sets.

Optimization for accuracy of quantification and depth of proteome coverage start from data acquisition. For example, duty cycles can incorporate multiple MS1 survey scans to increase the temporal frequency of sampling precursors and thus the probability of sampling close to the apex with reduced interference. Furthermore, duty cycle parameters can be algorithmically optimized to distribute ions evenly across isolation windows and thus reduce overcrowding of ions in some windows and the associated potential for increased interference. Such optimization can be performed with multiple tools, including with DO-MS^40,41^, which provides support for plexDIA.

Sequence propagation has long been fruitfully applied across different runs using a variety of software tools^42–46^. All these methods for matching between runs exploit retention time (RT) alignment between runs. However, even the best RT alignment between runs is likely to result in larger RT deviation than the one measured within a run. To precisely measure RT deviations within and between runs, we acquired plexDIA data with a duty cycle that included an MS1 survey scan before and after each MS2 scan. This resulted in frequent sampling of precursors, which supports good estimates of elution peak apexes, Fig. 3a. The data from these experiments indicate that indeed the RT deviations are smaller within plexDIA runs, as shown in Fig. 3b.

**Figure 3:**
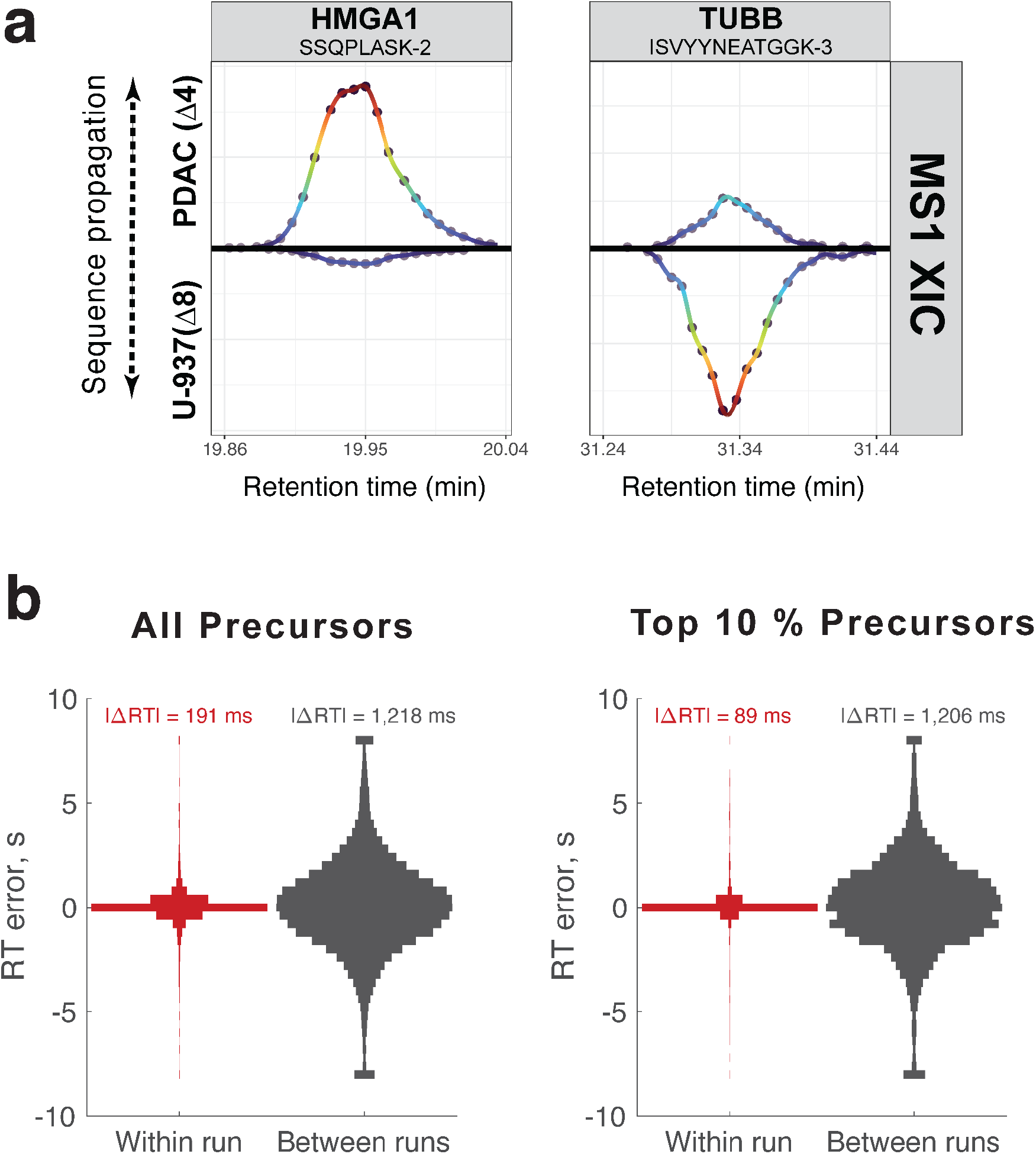
Precision of retention time estimates within and between plexDIA runs. **a**, Extracted ion current (XIC) for the precursors of two peptides quantified in U937 and PDAC cells. Circles correspond to measured inequalities and the curves are colored based on the intensity. **b**, The retention time (RT) deviations are estimated from triplicate injections of plexDIA samples composed of 100 cells of Melanoma, PDAC, and U-937 cells and analyzed with high frequency survey scans. Apex RTs within a run between channels comparing PDAC and U937 cells (“Within run”) are more similar than the aligned RTs (“Between runs”). The median absolute RT deviations (|ΔRT|) are indicated on top of each distribution in milliseconds (ms).

Even after RT alignment, the median RT difference for precursors across different runs, replicate injections of the same sample, is about 1.2 seconds both for all precursors and for the most abundant precursors, Fig. 3b. The median RT difference for precursors within a run is smaller, 0.19 seconds for all precursors and 0.09 seconds for the 10 % most abundant precursors, Fig. 3b. The smaller RT deviation for highly abundant precursors suggests that RT estimates for less abundant precursors are likely influenced by interferences. These data demonstrate that even for replicate injections runs one after another, plexDIA allows 6 – 13 fold higher precision in RT estimates between isotopologously labeled samples within the same run, as shown in Fig. 3. This gain is likely to be larger when comparing RT across diverse samples since variation in their protein composition and preparation may introduce further RT variability between runs.

Since the information content of RT estimates is directly proportional to their precision^47^, precise RT estimates increase the sensitivity and specificity of sequence propagation. Thus, the increased RT precision within plexDIA sets (Fig. 3) allows for more reliable sequence propagation than what can be achieved with methods matching RTs between runs. This benefit of precise RTs applies to both the precursors and all of their fragments and is thus compounded for plexDIA algorithms that propagate sequences using both precursors and their associated fragments. Therefore, we expect to see strategies for leveraging highly precise (and thus informative) RT estimates for enhancing the interpretation of plexDIA data.

### Isotopologous carriers for plexDIA

One such strategy suggested by Derks *et al*.^16^ is the introduction of isotopologous carriers that can facilitate the sequence identification in small samples, such as single cells. Specifically, Type 1 mass tags can be used to label both single cells and a carrier sample (composed of a small bulk sample and/or spiked in peptides of interest), and then peptide sequences identified in the carrier may be confidently propagated to the single cells. This strategy extends the ideas of isobaric carriers^20,38,48^ motivated us to develop plexDIA^36^. It appears promising, though it will encounter similar challenges as the isobaric carrier approach, such as the limited capacity of ion traps and detectors^20^. This capacity limitation is likely to be more pronounced for DIA analysis using isotopologous carriers because of the wider isolation windows used by DIA. Thus the development of isotopologous carriers can be informed by established principles for the use of carriers^4,20^, and isotopologous carriers compatible with accurate and precise quantification are likely to be smaller than isobaric carriers. Thus, the optimal carrier size and data acquisition methods should be rigorously benchmarked using mixed species experiments as with the development of plexDIA^16^.

## Applications

Increasing the throughput and depth of proteome coverage will empower many applications, especially those requiring many samples for estimating robust associations^8^ and protein covariation^49,50^. Furthermore, plexDIA can confer additional advantages. Below we highlight some examples, though we expect that the community will find many more.

### Increasing the stability of large-scale longitudinal studies

Proteomics is increasingly applied to clinical samples, and in many cases the samples may be collected and analyzed over a long period of time. If analyzed by LF-DIA, changes in peptide separation and MS acquisition (e.g., due to instrument drift) may be challenging to control and compensate for. Such undesired technical variability may be mitigated by including a common reference to all plexDIA sets analyzed. For example, a 6-plexDIA experiment may run five biological samples and a reference standard; the biological samples can be normalized to the unchanging reference to control for technical variability during data acquisition.

### Improved interpretation of missing values

The ability to confidently propagate peptide sequences within plexDIA sets allows detecting peptides present at levels that may not support confident sequence assignment with LF-DIA, even with MBR. This increased sensitivity of propagating sequences within a run effectively lowers the limit of detection and increases data completeness. As a direct consequence, it increases the confidence in interpreting missing values as corresponding to very low levels of peptide abundance, below the limit of detection, which can be useful for downstream data interpretation. For example, if an N-terminal peptide from a protein becomes undetectable in a plexDIA sample while all other peptides from the protein are detectable, we may infer that the abundance of the proteoform containing the N-terminal peptide has fallen below the limit of detection. This may be reflected in an alternative open reading frame or a post-translational modification, thus suggesting hypotheses for further investigation.

### Functional proteomic assays with plexDIA

In addition to mass tags for increasing sample throughput, the plexDIA framework can be extended to a wide array of functional assays using covalent protein modifications. One example includes footprinting methods^51^ using chemical labeling, such as dimethyl labeling of surface exposed lysines that allows quantifying protein conformations in live cells^52,53^. Different samples can be labeled with isotopically coded dimethyl tags and combined into a plexDIA set, whose analysis will take less time than analyzing each sample individually. Furthermore, it will benefit from increased data completeness and sensitivity arising from sequence propagation within a plexDIA set. Other example for functional assays that benefit from plexDIA include (i) quantifying regulatory proteolysis based on labeling free amine groups prior to protein digestion^54^, (ii) activity-based protein profiling (ABPP) employing isotopically coded molecular probes^55,56^, and (iii) plexDIA pulsed SILAC for measuring protein synthesis and degradation rates. Since these applications can involve efficient and specific binding of chemical probes or mass tags, the resulting modified peptides may be produced in stoichiometric amounts and thus quantifiable in very small samples, even in single cells^53^. Thus, such extensions of plexDIA hold the potential of extending the toolset of single-cell analysis to functional assays quantifying protein shapes and activities^4,49,57^.

## Methods

### Cell culture and sample preparation

Cells were cultured and prepared as previously described^16^. U-937 monocytes were cultured in RPMI 1640 Medium (Sigma-Aldrich, R8758) and supplemented with 10% FBS (Gibco, 10439016) and 1% penicillin–streptomycin (Gibco, 15140122). Pancreatic ductal adenocarcinoma (PDAC) cells (HPAF-II, ATCC CRL-1997) were cultured in EMEM (ATCC, 30-2003), and likewise supplemented with 10% FBS and 1% penicillin–streptomycin. Melanoma cells (WM989-A6-G3) (a kind gift from Arjun Raj at University of Pennsylvania) were cultured in TU2% media. All cells were grown at 37°C, harvested at a density of 10^6^ cells/mL, washed with sterile PBS, then resuspended to a concentration of 3×10^6^ cells/mL in pure LC-MS grade water, then stored at −80°C. Cell numbers were estimated and diluted to 100-cell samples as described in the SCoPE2 protocol^6^.

Cell suspensions were prepared for proteomic analysis by mPOP^58,59^. In short, the frozen samples were thawed, aliquoted to PCR tubes, heated at 90°C for 10 minutes in a thermal cycler, then digested with Trypsin Gold (Promega, V5280) at a 1:25 ratio of protease:substrate in the presence of 100 mM Triethylammonium bicarbonate (TEAB) and 0.2 units/μL Benzonase nuclease (Millipore, E1014) for 18 hours at 37°C. Melanoma, PDAC, and U-937 digests were labeled with mTRAQ-Δ0, mTRAQ-Δ4, and mTRAQ-Δ8 mass tags (SciEx, 4440015, 4427698 and 4427700), respectively, then pooled to form a plexDIA set.

### Data acquisition

For the purpose of assessing RT-deviations within a run and between runs (Fig. 3), we needed a data acquisition method that samples precursors with high frequency and thus allows for accurate estimation of elution peak apexes. We applied such a method to analyze triplicate plexDIA sets of 100-cell inputs of Melanoma, PDAC, and U-937 cells. Each plexDIA set was injected at 1 μL volumes with a Dionex UltiMate 3000 UHPLC to enable online nLC to separate peptides. Flow-rate was set to 200 nL/min, and the gradient was set as follows: 4% Buffer B until 2.5 minutes, ramp to 8% B by minute 3, ramp to 32% B by minute 33, ramp to 95% B by minute 34, hold at 95% B until minute 35, lower B buffer to 4% by minute 35.1, then hold at 4% B buffer until minute 60.

Mass spectrometry data acquired on a first-generation Q-Exactive Hybrid Quadrupole-Orbitrap with the following DIA duty cycle in positive ion mode using frequent survey scans, MS1 scans spanning the range 379-1401 m/z. The duty cycle was: 1 survey scan, 1 MS2 (380-460 m/z), 1 survey scan, 1 MS2 (460-540 m/z), 1 survey scan, 1 MS2 (540-620 m/z), 1 survey scan, 1 MS2 (620-740 m/z), 1 survey scan, 1 MS2 (740-980 m/z), 1 survey scan, 1 MS2 (980-1400 m/z). Each MS1 was performed at 70k resolving power, 240 ms max fill time, and 3×10^6^ AGC max. Each MS2 was performed at 35k resolving power, 110 ms fill time, and 3×10^6^ AGC max, and 27 NCE with default charge of 2. This method enables high temporal resolution of MS1 features.

### Data analysis

#### RT-deviations within a run and between runs

Raw files from triplicate plexDIA sets of 100-cell Melanoma, PDAC, and U-937 cells were searched by DIA-NN^13^ version 1.8.1 with the following commands: {fixed-mod mTRAQ, 140.0949630177, nK}, {channels mTRAQ,0,nK,0:0; mTRAQ,4,nK,4.0070994:4.0070994; mTRAQ,8,nK,8.0141988132:8.0141988132}, {peak-translation}, {original-mods}, {report-lib-info}, {ms1-isotope-quant}. This search used the spectral library that was previously generated from 100-cell plexDIA runs of Melanoma, PDAC, and U-937 cells^16^.

Peptide-like features were extracted from the raw files by processing with Dinosaur^60^. Precursors which were quantified as reported by DIA-NN in all three channels were mapped to the corresponding features +/− 5 ppm and with the apex RT falling within the elution start and stop RTs as reported by DIA-NN. The ‘within run’ condition compared apex RTs of PDAC cells to U-937 cells as reported by Dinosaur as they co-eluted within a run. The ‘between run’ condition subtracted the ‘Predicted.RT’ column output by DIA-NN to the apex RT reported by Dinosaur. This was performed for all precursors and for the top 10% most abundant precursors averaged between U-937 and PDAC channels across triplicates.

#### Benchmarking accuracy of translated quantitation

The mixed-species plexDIA data used in Fig. 2 to quantify the accuracy of protein quantification was previously generated^16^ and are available at MassIVE: [MSV000089093](https://massive.ucsd.edu/ProteoSAFe/dataset.jsp?task=ae918c7ce5a94a4abd2c6b54a3806c9e). The errors were estimated as the difference between the measured and mixing proteome ratios^16^. DIA-NN reports which have columns ‘MS1.Area’, ‘Ms1.Translated’, ‘Precursor.Quantity’, and ‘Precursor.Translated’ correspond to MS1 or MS2-level quant that is either translated or not-translated. These values were used to compute the empirically observed precursor ratios from the DIA-NN report of a single raw file, ‘wJD804’. Empirically observed ratios and expected ratios were log transformed, then subtracted from each other, then the absolute value was plotted as boxplots to display the errors of precursor quantitation.

## Data availability

Raw files, spectral library, and DIA-NN and Dinosaur outputs can be found at MassIVE MSV000090650 and https://scp.slavovlab.net/plexDIA.

## Code availability

Code used for data analysis can be found at: https://github.com/SlavovLab/plexDIA_perspective.

## Acknowledgements

This work was supported by an Allen Distinguished Investigator award through The Paul G. Allen Frontiers Group to NS, a Seed Networks Award from CZI CZF2019-002424 to NS, and an R01 by NIGMS 5R01GM144967 to NS.

